# Detecting the state of drowsiness induced by propofol by spatial filter and machine learning algorithm on EEG recording

**DOI:** 10.1101/2021.07.12.452077

**Authors:** Zhibin Zhou, Ramesh Srinivasan

## Abstract

To accurately measure the depth of anesthesia has been a challenge for both anesthesiologists and engineers who work on developing tools of measurements. This study aims to use a machine-learning algorithm to predict the drowsy state, a transitional depth of sedation during propofol anesthesia. The data used in this study were scalp EEG (electroencephalogram) recordings selected from the University of Cambridge Repository. Raw EEG recordings were preprocessed into power spectrum matrices one second per sample. A total of 170 samples (110 awake samples and 60 drowsy samples) were used. A CNN (Convolutional Neural Network) for the MNIST (Modified National Institute of Standards and Technology) dataset was applied on these EEG power spectrum matrices. Due to the small dataset volume, Leave-One-Out cross-validation was used to train the data. Results of the training accuracy reached 99.69%. And test accuracy averaged 96.47%. Overall, the model is able to predict the state of drowsiness during propofol anesthesia. This provides the potential to develop EEG monitoring devices with closed-loop feedback of such a machine learning algorithm that controls the titration of the dosage of anesthetic administration and the depth of anesthesia.

## 1 Introduction

### 1.1 Motivation & specific questions

Propofol is the most commonly used intravenous anesthetic drug today. Millions of people in the world take surgeries and minor procedures under propofol anesthesia every day. In larger dosages, propofol induces unconsciousness (drug-induced coma) so that major surgeries can be performed. However, larger doses come with higher risks. Larger propofol dosage suppresses autonomic protective nerve reflects, meaning the risk of aspiration, respiratory suppression, hemodynamic instability, hypothermia, etc[1]. In lower dosage, it induces a drowsy state so that patients can tolerate minor procedures without much discomfort, meanwhile with benefits such as preservation of autonomic respiratory, protective gag reflex, and stable hemodynamics [2]. As a result, it is preferred to maintain patients under such a state of drowsiness, so that small procedures such as colonoscopy can be well tolerated and the risk of anesthesia complications can be minimized[3].

Accurately measuring the neural correlates of consciousness is a challenge for neuroscience, let alone the depth of anesthesia. One main feature of the EEG under anesthesia is the domination of slow-wave oscillation [4]. Slow-wave activities will increase after the level of consciousness decreases in the drowsy state[5]. The other important feature is the shifting of alpha wave oscillations from the occipital lobe to the frontal lobe, which is prominent under propofol anesthesia[6]. The level of alpha activities seems to play an important role in the responses under the influences of anesthesia [7].

Our aim is to test whether a convolutional neural network will be able to detect the change of EEG features under anesthesia, especially on the features in frequencies ranging from the slow-wave domain to the alpha band domain.

### 1.2 The selection of training and testing dataset & preprocessing

The training and testing samples were selected from the wake states and drowsy states in patients showing a higher percentage of correct responses and lower correct responses in a simple behavioral task involving fast discrimination between two possible auditory stimuli.

In a pilot analysis, we used PCA (Principle Component Analysis)[8] and LDA (Linear Discrimination Analysis)[9] to separate the two states of consciousness (the awake and drowsy states under propofol anesthesia). We had found that the alpha components in the data have a distinct spatial pattern that separates the two states. As a result, the training and testing dataset selected for machine learning was mainly power spectrum matrices focusing on the alpha power distribution.

We’ve used Fast Fourier Transformation to extra alpha components from the original EEG data. We combined two types of analysis, the PCA (Principal Component Analysis)[8] and LDA (Linear Discrimination Analysis)[9] to source the principal components of the alpha activities and to maximally separate the two states of consciousness, the awake and the drowsy state under anesthesia. PCA and LDA will be integrated together as a Spatial Filter to identify the spatial pattern responsible for the separation of two conscious states.

The hypothesis is, the spatial differences in the distribution of alpha power combined with that of delta power will be detected by the convolutional neural network model, which can be predicted when such a drowsy state of power spectrum occurs during propofol anesthesia. The machine learning algorithm can further serve as a core function in the development of a closed-loop feedback system on controlling the dosage of anesthetic administration, enhancing existing EEG monitoring devices to help physicians titrate anesthetic dosage to achieve the optimal state of sedation.

## 2 Methods

### Source of the data

University of Cambridge Repository [7] (https://www.repository.cam.ac.uk/handle/1810/252736)

### Data type

Dense array EEG during the awake and drowsy state under propofol anesthesia. A total of 12 subjects were included in this study.

### Experimental conditions

Subjects were gradually induced into different sedation levels according to TCI (Target Control Infusion) plasma propofol concentrations. EEG was recorded during Baseline (the awake state), Mild Sedation, Moderate Sedation (the drowsy state), and Recovery. There are 91 channels. The sampling frequency is 250. Each epoch of data is 7 minutes long, both in the awake and drowsy states.

### Time-frequency Analysis

The original EEG data were segmented into one-second epochs. Fast Fourier Transformations were performed to extra the power of alpha frequency at 11Hz for each epoch. This creates two alpha power matrices for both the awake state and the drowsy state over all 91 channels.

### Separation of the alpha band distribution

Using Principal Component Analysis, the two alpha power matrices can be transformed into 91 principal components. The first ten principal components were selected for further analysis. For Linear Discrimination Analysis, these two alpha power matrices were submitted to our customized LDA algorithm with 10-fold cross-validation. We then averaged the 10 models together, since each model is based on 90% of the data in the alpha matrices. The choice of the number of components in PCA and the consistency of the results from our classifier were tested with the Akaike information criterion (AIC) and the error rate. These processing steps were implemented using customized MATLAB scripts. Two random subject epochs were selected as the test signal. The results included 12 subjects (6 subjects in both awake and drowsy groups).

### Preparation of the training and testing dataset

EEG power spectrum matrices were produced as the training and testing data samples using customized Matlab code in the Human Neuroscience Lab [http://hnl.ss.uci.edu/]. Frequencies higher than 12 Hz were removed from the power spectrums due to the lack of distinctive neural activities features. Features of the power spectrum in the delta range and alpha range were kept to be the main focus for machine learning. The training dataset consisted of 110 samples of matrices with the size of 12 by 91. The testing dataset contained 60 samples of such matrices. The data structure of the targets are vectors created with either one or zero. One for indicating the drowsy states. Zero for the awake states.

### Implementation of the machine learning algorithm

A CNN (Convolutional Neural Network) for the MNIST (Modified National Institute of Standards and Technology) dataset was adapted and applied on these EEG power spectrum matrices. Due to the small dataset volume, a Leave-One-Out cross-validator [10] was used to train the data. Data of the matrices and the code for this adapted neural network were stored in a public GitHub Repository (https://github.com/zhibinz2/Neural-Networks-and-Machine-Learning ). The neural network model was trained online using Google Colab.

## 3 Results

### Time-frequency and separation of the alpha band distribution

A total of 2580 epochs (trials) from 12 subjects in both the awake (responsive) and the drowsy group (6 subjects in each state) were extracted. Alpha power matrices for each state were plotted against the 91 channels on heat maps as in Figure 1. More alpha power was seen in the awake (responsive) group, appearing across more channels. The alpha power was not evenly distributed. Significant spatial distribution differences were observed.

**Figure 1:**
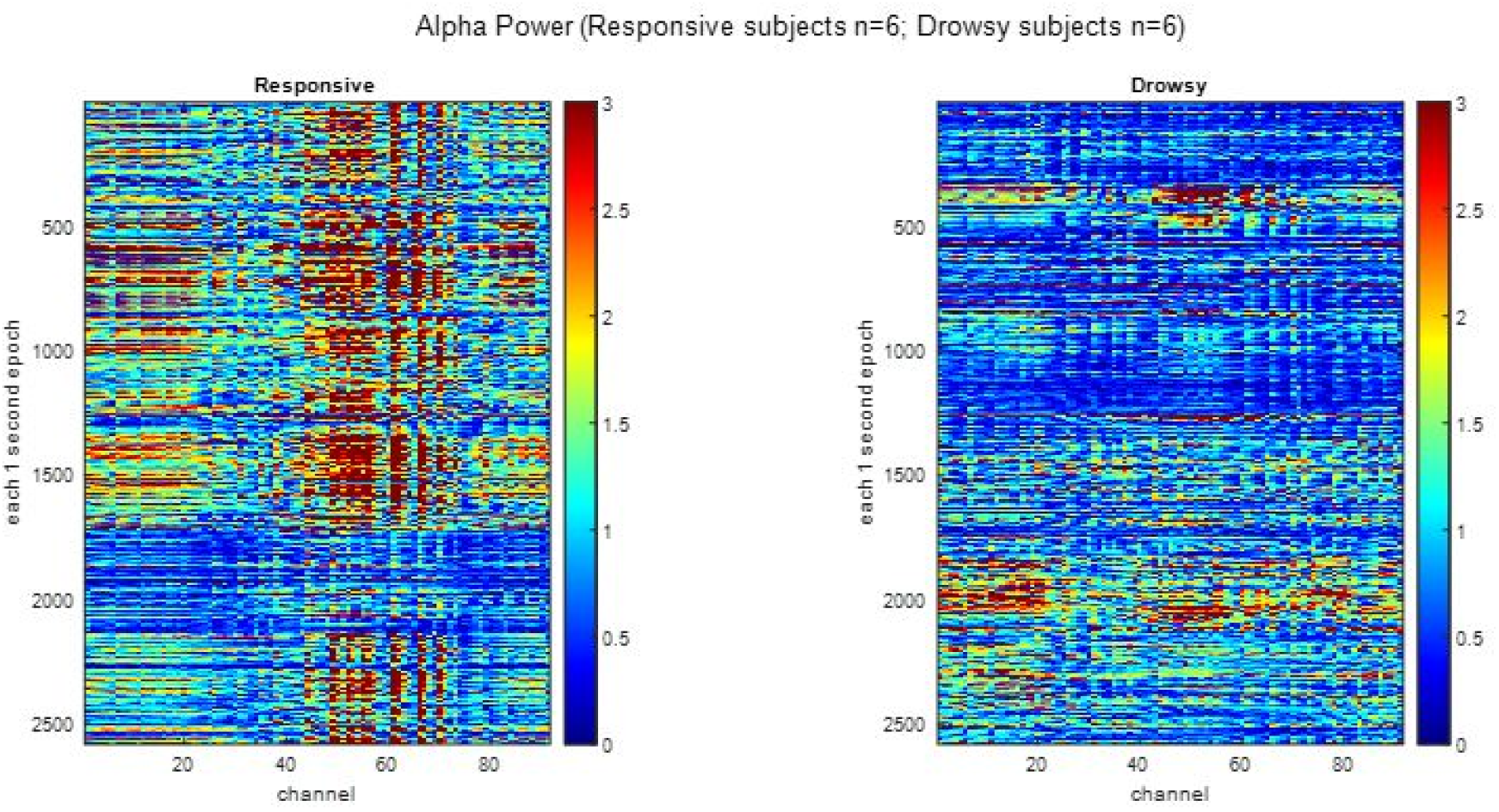
Sample Figure Caption

The choice of the number of components is a free parameter. The results of the principal component analysis are shown in Figure 2. The percentage accounted for the total variance for the 91 principal components was plotted. The first 10 to 20 principal components account for the most variance. We used AIC to examine the model order. The first 10 principal components were tested as shown in Figure 3.

**Figure 2.**
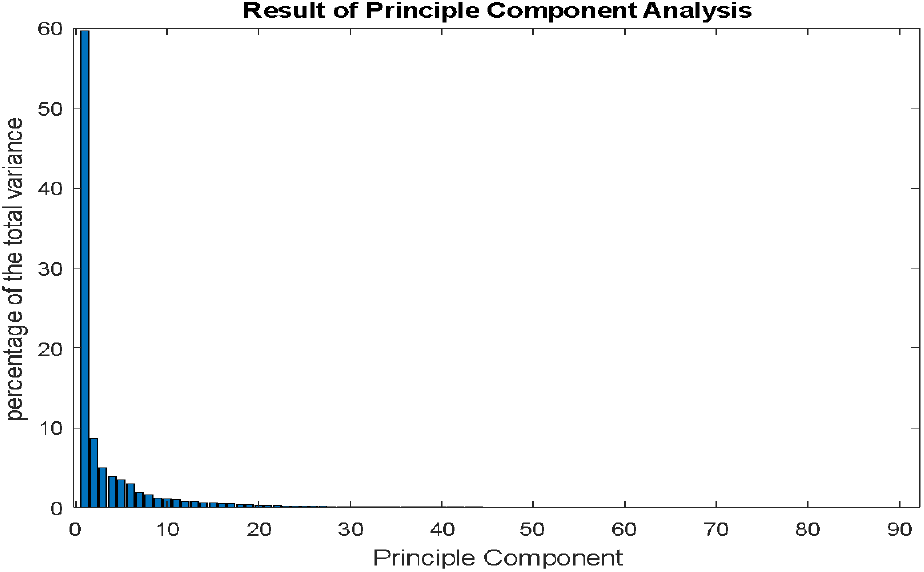
Result of Principle Component Analysis

**Figure 3.**
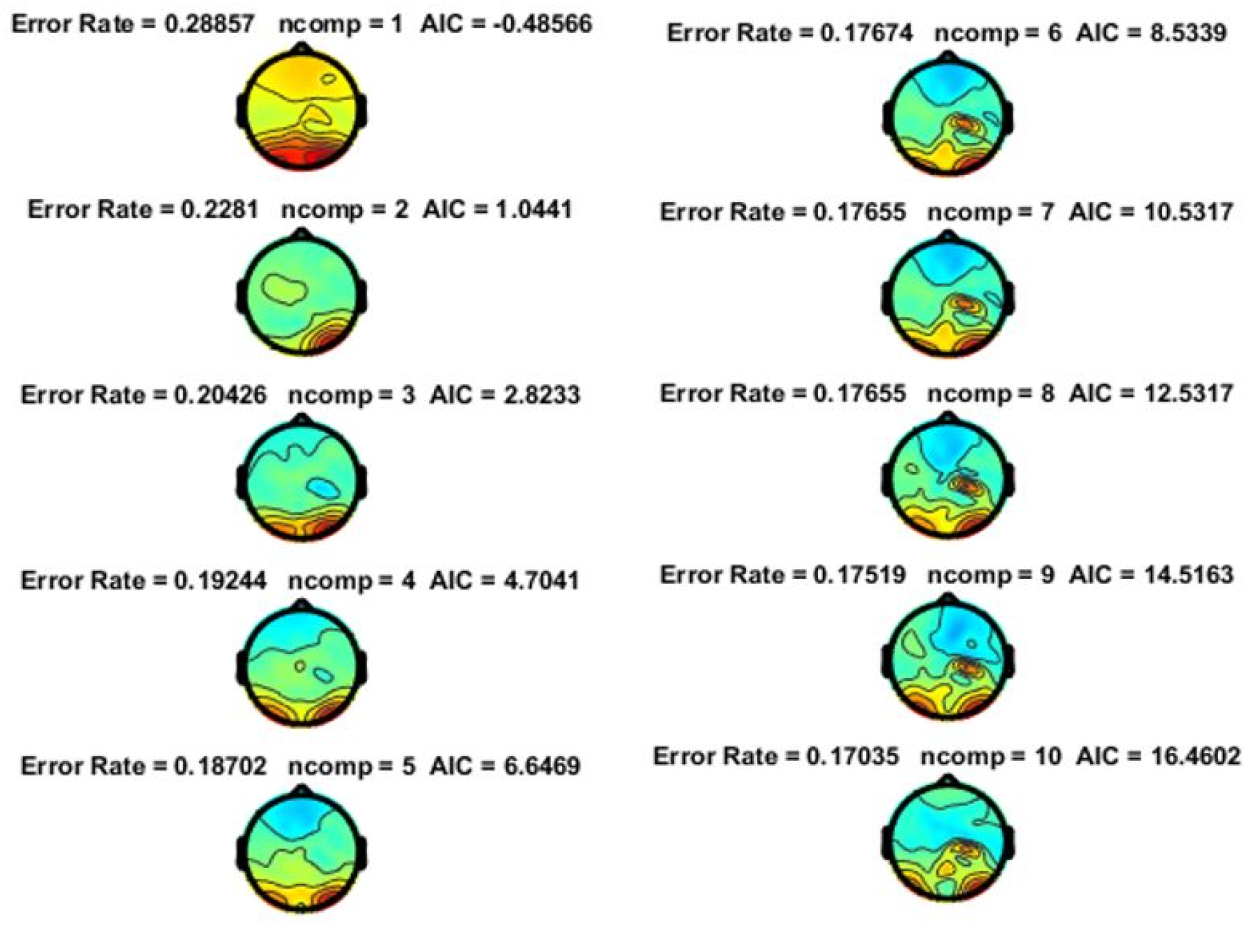
First 10 Principal Components

LDA was then done on the PCA data to find out the direction to separate the data. The topo plot of the spatial filter (Figure 4.) shows how much weight to put on each electrode to rotate the alpha power matrices and get a new set of weighted average variables which best separate the alpha pattern.

**Figure 4.**
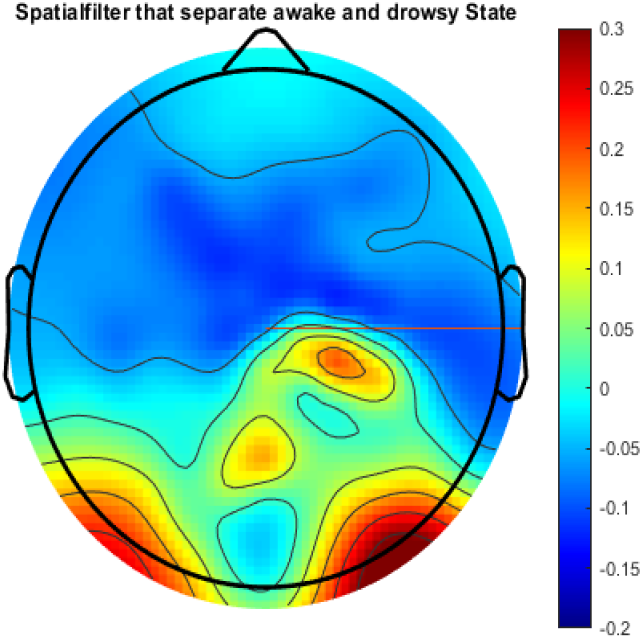
Spatial Filter That Separate Awake And Drowsy State

The projected data (Figure 5.) is the coordinate of each sample in a new coordinate system. It is a weighted average of the original data. The weights are given by the spatial filter. The spatial filter reduced the data into one dimension that best separated the two states as shown in Figure 4. The two conditions are significantly different with alpha power in the awake state mostly positive and the alpha power in the drowsy state mostly negative.

**Figure 5.**
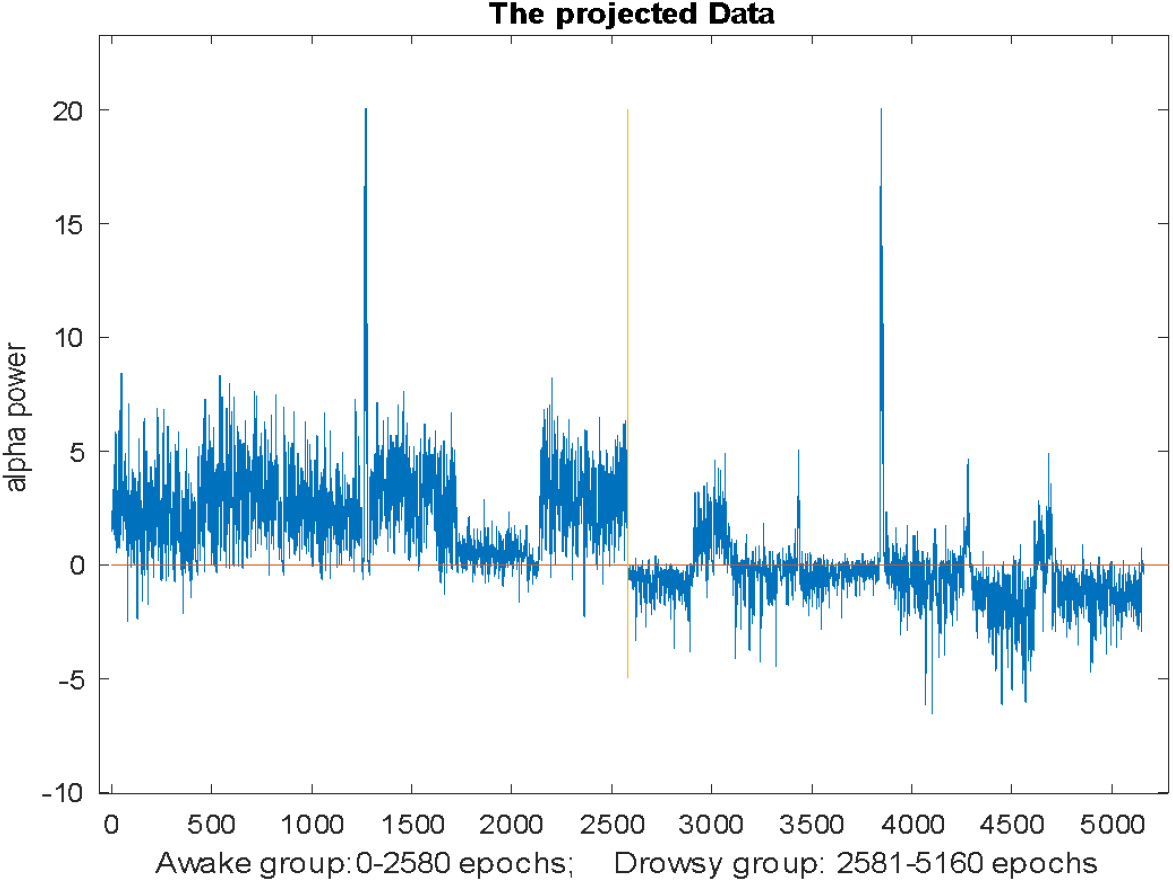
The Projected Data (Awake group: 0-2580 epoch; Drowsy group: 2581-5160 epochs)

The sample average of the EEG power spectrum focusing the delta band to alpha band distribution in the awake states and drowsy states are shown in Figure 6. Each Leave-One-Out training and testing cycle consists of 60 epochs. The average increment of training accuracy and testing accuracy over each epoch in one training cycle of the convolutional neural network model are demonstrated in Figure 7.

**Figure 6.**
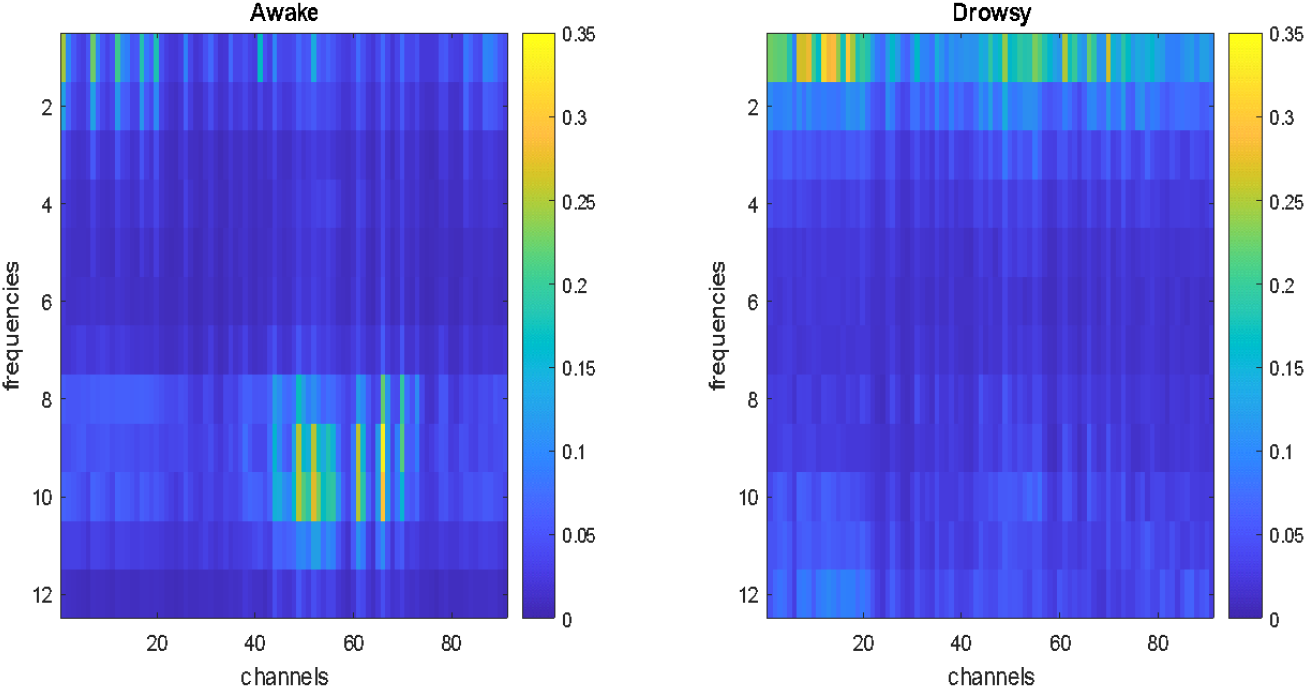
Average power spectrum of the awake state and drowsy state

**Figure 7.**
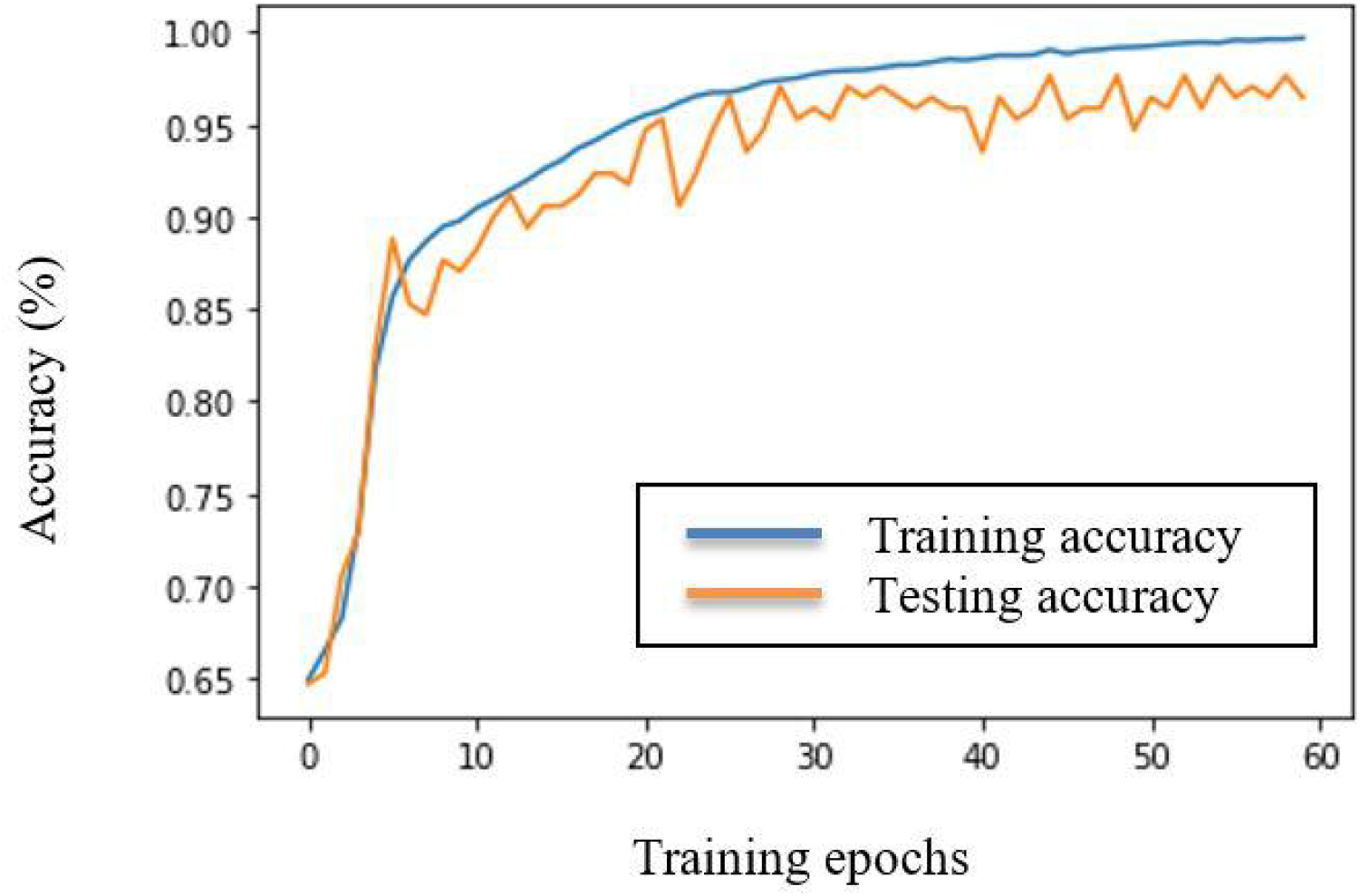
The average increment of training accuracy and testing accuracy over each epoch in one training cycle of the Leave-One-Out cross-validation (Training accuracy at epoch 60 reaches 99.69%; Testing accuracy at epoch 60 reaches 96.47%)

## 5 Discussion

The results of the pilot analysis support the hypothesis that a spatial pattern for the difference in alpha power distribution between the awake state and drowsy states under propofol anesthesia exists (Figure 3). This spatial pattern is able to separate the awake state and drowsy state effectively, which provides a solid foundation for the successful implementation of a neural network to predict such differences in the spatial distribution of the power spectrum. The reason that PCA and LDA are chosen is that that, on one hand, PCA can reduce the dimensionality of the original dataset, increasing the interpretability of the principal alpha components and at the same time reducing the effect of noise. On the other hand, LDA is widely used in pattern recognition and machine learning. It can be employed to find a linear combination of features in the alpha activities that characterize and maximally separate the two conscious states. The combination of PCA and LDA will serve as a spatial classifier for our purpose. Although there is a trade-off between a lower AIC (which is based on the chosen number of principal components) and a higher error rate (which relies on the LDA classifier) as shown in Figure 3. The spatial pattern of the classifier (as shown in Figure 4) for the first 10 principal components are very much consistent (as shown in Figure 5) with the most weight putting mainly on the occipital lobe and far less weight on the frontal lobe. The broader impact for this study is when applying this spatial filter to the monitoring data in real time, it has the potential to be developed into more advanced technologies for the clinician. So that the first-line health care workers can better maintain the safety and comfort for patients who need sedation or anesthesia for minor procedures. Future work for us is that further reduce the error rate in the spatial filter, which is around 12-17 % so far.

Testing accuracy of 96.47% is satisfactory so far, as the contemporary measurement using a combination of EEG and hemodynamic variables indicates the highest overall accuracy to predict the depth of anesthesia is 89.4%[10]. We will continue to seek better and refiner the machine learning models further as we include more samples in the dataset pool for training and testing. It will significantly improve perioperative health care quality to develop a machine-learning paradigm that can accurately detect the achievement of the drowsy states as the new monitoring data come in. A paradigm such as this can be applied in clinical EEG monitoring in real-time, guiding anesthesiologists and critical care physicians to determine the optimal dosage of anesthetic for small diagnostic and surgical procedures.

## Acknowledgments

We thank Professor Emre Neftci at the University of California, Irvine for his constructive advice for the implementation of this convolutional neural network and for adopting the Leave-One-Out cross-validation to make up for dataset sample volume. Special Thanks to the Department of Clinical Neurosciences, Division of Anaesthesia, and Department of Psychology at the University of Cambridge for providing this data repository. Customized codes used in preprocessing the EEG data in this study come from previous work done under Professor Ramesh Srinivasan’s Human Neuroscience Lab at the University of California, Irvine.

